# MicroRNA-375 is induced during astrocyte-to-neuron reprogramming and promotes survival of reprogrammed neurons when overexpressed

**DOI:** 10.1101/2023.07.10.548401

**Authors:** Xuanyu Chen, Ivan Sokirniy, Xin Wang, Mei Jiang, Natalie Mseis-Jackson, Christine Williams, Kristopher Mayes, Na Jiang, Brendan Puls, Quansheng Du, Yang Shi, Hedong Li

## Abstract

While astrocyte-to-neuron (AtN) reprogramming holds great promise in regenerative medicine, the molecular mechanisms that govern this unique biological process remain elusive. MicroRNAs (miRNAs), as post-transcriptional regulators of gene expression, play crucial roles during development and under various pathological conditions. To understand the function of miRNAs during AtN reprogramming process, we performed RNA-seq of both mRNAs and miRNAs on human astrocyte (HA) cultures upon NeuroD1 overexpression. Bioinformatics analyses showed that NeuroD1 not only activates essential neuronal genes to initiate reprogramming process but also induces miRNA changes in HA. Among the upregulated miRNAs, we identified miR-375 and its targets, *neuronal ELAVL* genes (*nELAVLs*), which encode a family of RNA-binding proteins and are also upregulated by NeuroD1. We further showed that manipulating miR-375 level regulates *nELAVLs* expression during NeuroD1-mediated reprogramming. Interestingly, miR-375/*nELAVLs* are also induced by reprogramming factors Neurog2 and ASCL1 in HA suggesting a conserved function to neuronal reprogramming, and by NeuroD1 in the mouse astrocyte culture and spinal cord. Functionally, we showed that miR-375 overexpression improves NeuroD1-mediated reprogramming efficiency by promoting cell survival at early stages in HA even in cultures treated with the chemotherapy drug Cisplatin. Moreover, miR-375 overexpression doesn’t appear to compromise maturation of the reprogrammed neurons in long term HA cultures. Lastly, overexpression of miR-375-refractory ELAVL4 induces apoptosis and reverses the cell survival-promoting effect of miR-375 during AtN reprogramming. Together, we demonstrate a neuro-protective role of miR-375 during NeuroD1-mediated AtN reprogramming and suggest a strategy of combinatory overexpression of NeuroD1 and miR-375 for improving neuronal reprogramming efficiency.

## INTRODUCTION

Despite controversies around the validation of reprogrammed neurons *in vivo* (Wang et al., 2021a; Xiang et al., 2021), glia-to-neuron reprogramming has been successfully demonstrated in several laboratories (Guo et al., 2014; Heinrich et al., 2014; Qian et al., 2020; Su et al., 2014). This has been achieved mainly through overexpression of neurogenic transcription factors, either alone or in combination, in various neurological disease or injury models (Chen and Li, 2022; Li and Chen, 2016; Tai et al., 2020). One such factor, NeuroD1, has been shown to convert astrocytes, NG2 glia, and microglia into functional neurons both *in vitro* and *in vivo* (Brulet et al., 2017; Guo et al., 2014; Matsuda et al., 2019; Puls et al., 2020). Our previous study also demonstrated that ectopic expression of NeuroD1 reprograms reactive astrocytes to functional neurons in the injured spinal cord (Puls et al., 2020). NeuroD1 is widely expressed in the developing central nervous system (CNS) and is critical to neuronal differentiation (Miyata et al., 1999). NeuroD1 is also a “pioneer” transcription factor (Iwafuchi-Doi and Zaret, 2014), and reprograms the chromatin landscape to elicit neuronal programming in embryonic stem (ES) cells (Pataskar et al., 2016) and microglia (Matsuda et al., 2019) when overexpressed. Exogenously expressed NeuroD1 can activate expression of downstream target genes such as Hes6 and NeuroD4 (Pataskar et al., 2016), which may serve as important effectors for neuronal conversion. While neuronal reprogramming has become a feasible approach for neuroregeneration, our understanding of the molecular mechanisms that govern this unique biological process is still incomplete.

MicroRNAs (miRNAs) are small non-coding RNAs that regulate gene expression post-transcriptionally (Bartel, 2004), and are crucial to the differentiation of neural cell types during CNS development (Rajman and Schratt, 2017). Previous work from our own lab and others demonstrated that miRNAs are important to cellular differentiation in the developing mouse forebrain (Davis et al., 2008; McLoughlin et al., 2012; Nowakowski et al., 2011; Zhang et al., 2015) and cerebellum (Kuang et al., 2012; Liu et al., 2017; Tao et al., 2011; Zindy et al., 2015), and are indispensable to reactive astrogliosis during spinal cord injury (Hong et al., 2014). However, the function of miRNAs during neuronal reprogramming has not been systemically investigated.

In this study, we aim to decipher miRNA function during NeuroD1-mediated neuronal conversion using human astrocyte (HA) culture as a model. Our RNA-seq analyses showed that NeuroD1 overexpression induces drastic upregulation of two miRNAs, miR-375 and miR-124. We experimentally demonstrated that miR-375 modulates expression level of *neuronal ELAVL* genes (*nELAVLs*), which encode RNA-binding proteins and are also upregulated by NeuroD1. Interestingly, miR-375/*nELAVLs* are also induced by reprogramming factors Neurog2 and ASCL1 in HA, and by NeuroD1 in the mouse astrocyte culture and spinal cord. Furthermore, overexpression of miR-375 by a retrovirus promotes cell survival during NeuroD1-mediated astrocyte-to-neuron (AtN) reprogramming at early stages without compromising maturation of reprogrammed neurons in HA cultures. Thus, our data indicate that miR-375 facilitates NeuroD1-mediated reprogramming by modulating expression level of target genes including *nELAVLs*, and that combinatory overexpression of NeuroD1 and miR-375 may represent an improved reprogramming strategy with a greater translational value.

## EXPERIMENTAL PROCEDURES

### Animal use

Wild-type C57BL/6 mice (2–4 months old) were used for AAV injection experiments. Mice were housed in a 12 h light/dark cycle and supplied with sufficient food and water. All animal use and studies were approved by the Institutional Animal Care and Use Committee (IACUC) of Augusta University. All procedures were carried out in accordance with the approved protocols and guidelines of National Institute of Health (NIH).

### Virus production

Retroviruses expressing NeuroD1-GFP and GFP control have been previously described (Guo et al., 2014). For NeuroD1-RFP, NeuroD1-coding sequence was subcloned into pCAG-Neurog2-IRES-DsRed vector (Heinrich et al., 2010) to replace Neurog2 by PCR-based strategy. For miR-375-GFP, a 450bp genomic sequence containing human miR-375 mature sequence was cloned into NeuroD1-GFP vector by replacing NeuroD1 at PmeI and NotI sites. For ELAVL2-GFP and ELAVL4-GFP, human gene coding sequences were cloned into NeuroD1-GFP vector by replacing NeuroD1 at PmeI and NotI sites. Lenti-miR-375-decoy was a gift from Dr. Brian Brown (Addgene plasmid # 46617).

Retrovirus packaging was performed as described (Guo et al., 2014). For Retro-miR-375-GFP packaging, a plasmid containing human Dicer shRNA [pSicoR human Dicer1, a gift from Tyler Jacks (Addgene plasmid # 14763)] was included to increase virus yield (Chang et al., 2013; Liu et al., 2010). For lentivirus packaging, the procedure was similar to that of retrovirus except that the packaging plasmids psPAX2 and VSV-G were used. Virus-containing culture media were centrifuged and filtered to remove cell debris, then aliquoted and stored at −80°C before use.

Virus media were routinely assayed for titer by infecting HEK293T cells. Our retrovirus and lentivirus titers are usually at 10^7^ genomic copies (GC)/ml. AAV viral particles were produced using the established procedure (Challis et al., 2019). HEK293T cells were transfected with three plasmids: a pAAV of interest (pAAV-Flex-GFP, pAAV-Flex-ND1-GFP, pAAV-GFAP-Cre), pAAV2/5 (Addgene Plasmid #104964), and pHelper (Cell Biolab) vector with 1:4:2 ratio, respectively. Medium containing viral particles was collected at 72 and 120 hours after transfection. After centrifuging at 2,000 × g for 15 min, the supernatant was precipitated in 40% polyethylene glycol (PEG) in 2.5 M NaCl. Following centrifuging this PEG-media mixture at 4,000 × g for 30 min, we combined the PEG pellets with the cell pellets in lysis buffer (500 mM NaCl, 40 mM tris, 10 mM MgCl2, and salt-active nuclease). The resulting lysates were extracted from an iodixanol step gradient following ultracentrifugation of 350,000 × g for 2.5 h. Purified AAVs were concentrated with Amicon Ultra-15 centrifugal filter devices. The AAV titers were determined by quantitative PCR (qPCR) as described (Aurnhammer et al., 2012). AAV mixtures with a titer of 1 × 10^12^ GC/ml were used for spinal cord injection experiments.

### Human astrocyte culture, virus infection, and Cisplatin treatment

The procedures for HA cultures (HA1800, ScienCell) and medium change after virus infection (Guo et al., 2014) were previously described. In some experiments, spinfection was performed to increase infection efficiency (Lu et al., 2021). Briefly, virus-containing media were mixed with polybrene (4 µg/ml) before being applied onto HA cultures. The resulting culture plates were centrifuged at 1,000 × g for 45 min at room temperature and then cultured in CO_2_ incubator at 37°C. At one day post virus infection, cultures were subject to medium change from HA culture medium (HAM) to neuron differentiation medium (NDM) (Wang et al., 2021b). For long-term cultures, NDM was changed every four days. Brain-derived neurotrophic factor (BDNF, 20 ng/ml, Invitrogen) was supplemented to support survival of reprogrammed neurons (Guo et al., 2014). Treatment of Cisplatin (Sigma-Aldrich) was done at indicated concentrations for 24 h in HA cultures before fixation and immunostaining.

### RNA-seq and bioinformatics analyses

Total RNAs were extracted from cultured cells by using TRIzol® reagent (Invitrogen). The integrity of the total RNAs was assessed using the Bioanalyzer 2100 (Agilent). We then made a uniquely indexed library from each sample using the Illumina TruSeq® Stranded mRNA library kit (Illumina). We also separately made a uniquely indexed miRNA library from each sample using the Illumina Small RNA library kit (Illumina). We made an equimolar pool of the mRNA libraries and a separate equimolar pool of the miRNA libraries, and then mixed the two library pools together at a ratio of 85 parts mRNA library pool: 15 parts miRNA library pool. A 75-nt single-read sequencing run on this combined pool was performed on the Illumina MiSeq® sequencer (Illumina) at Penn State Genomics Core Facility. The reads were divided up as ∼25 million reads per sample for the mRNA libraries and ∼5 million reads per sample for the miRNA libraries.

For both mRNA and miRNA sequencing, FastQC (v0.11.9) was used to check the quality and adaptor contents of the reads. For mRNA sequencing, adaptor contents were low, and no trimming was needed. The Fastq read files were quantified using kallisto (v0.46.1) with the transcriptome indices built on the cDNA FASTA files from human genome assembly GRCh38. Genes with lower than 10 total read counts were filtered for subsequent differential expression analysis. Differential expression analysis was performed using Deseq2 (v1.32.0) by comparing different samples, and genes with fold changes > 2 and p-value < 0.05 were called as differentially expressed. Gene Ontology (GO) Consortium and KEGG pathway databases were used to perform GO and pathway enrichment analysis in DAVID (https://david.ncifcrf.gov). Gene set enrichment analysis (GSEA) was used to check the enrichment of cell type signatures from cluster markers identified in single-cell sequencing studies of human tissue (C8). Venn diagram analyses were performed in InteractiVenn (http://www.interactivenn.net/). Gene network analysis was performed using STRING database (http://string-db.org). For miRNA sequencing, the reads were trimmed using Trim Galore! 9 (v0.6.5) for adapter sequences and any trimmed reads greater than 30 bp were filtered. The remaining reads were aligned to the genomic DNA sequence from human genome assembly GRCh38 using Bowtie2 (v2.4.3), and the aligned reads for each miRNA were quantified with featureCounts in the Subread package (v2.0.3) according to the sequences of all miRNA hairpins from *miRbase*. MiRNAs with lower than 5 total read counts were filtered for subsequent differential expression analysis. Differential expression analysis was performed using Deseq2 (v1.32.0) by comparing different samples, and miRNAs with fold changes > 1.5 and p-value < 0.05 were called as differentially expressed.

### Laminectomy, spinal cord injury, and stereotaxic viral injection

Mice were anesthetized using a SomnoFlo™ Low-flow electronic vaporizer connected with isoflurane. A laminectomy and stab injury were then performed as previously described (Puls et al., 2020). Briefly, after exposing the dorsal surface of the spinal cord at the T11–T12 vertebrae, the longitudinal stab injury was conducted with a 26-gauge needle with 0.8 mm in depth and 2 mm in length. The lesion site is 0.3 mm lateral to the central artery. By using a 50-μl Hamilton syringe with a 34-gauge injection needle, 1 μl of AAVs was immediately injected at the proximal and distal lesion sites at a depth of 0.7 mm and at a rate of 0.1 μl/min and the needle was gradually moved up to a depth of 0.3 mm during the injection. The injection needle was kept in place for 5 min after injection to prevent drawing out the virus during withdrawal and then slowly withdrawn. The mice were kept on a heating pad and treated with carprofen (5 mg/kg) for pain relief via subcutaneous injection.

### Mouse astrocyte isolation and culturing

For mouse astrocyte culture, postnatal (P1–P3) mouse cortical tissue was dissociated and plated onto 25 cm2 flasks (Guo et al., 2014). Cells were cultured for 7 days, and flasks were rigorously shaken to remove small non-astrocytic cells. After reaching confluence, astrocytes were passaged and plated on poly-D-lysine (Sigma) coated coverslips before being infected by retroviruses the following day. One day later, the medium was changed to NDM for reprogramming. Astrocyte culture medium contained DMEM/F12 (Gibco), 10% FBS (Gibco), penicillin/streptomycin (Gibco), 3.5 mM glucose (Sigma), and supplemented with B27 (Gibco), 10 ng/ml EGF and FGF2 (Invitrogen).

### Fluorescence activated cell sorting (FACS)

Three days after infecting HA with ND1-RFP and miR-375-GFP retroviruses, cultures were passaged and resuspended in medium containing 2% FBS. To exclude non-viable cells, 1 μl of 7-AAD staining solution (Enzo, 1 mg/ml) was added to cell suspension prior to sorting. GFP and RFP cell sorting was performed on the MoFloXDP 2-laser, 7-color cell sorter using Summit software (Beckman Coulter Life Sciences). The sorted RFP+ only and RFP+/GFP+ cell populations were collected and replated on coverslips with pre-seeded HA cells in NDM. The sorted cells were maintained in culture and fixed at the indicated time for immunocytochemistry.

### Immunocytochemistry and immunohistochemistry

Immunocytochemistry was carried out as previously described (Guo et al., 2014). Briefly, fixed cell cultures were incubated with monoclonal antibodies against ELAVL4 (mouse IgG, 1:200, Santa Cruz), NeuroD1 (mouse IgG, 1:200, Abcam), GFP (rat IgG2a, 1:400, BioLegend), and NeuN (mouse IgG, 1:400, Millipore); polyclonal antibodies against GFP (chicken IgY, 1:400, Aves), mCherry/RFP (rabbit IgG, 1:500, Abcam), mCherry/RFP (chicken IgY, 1:400, Aves), Map2 (rabbit IgG, 1:400, Abcam), DCX (rabbit IgG, 1:500, Abcam), ELAVL2 (rabbit IgG, 1:100, Santa Cruz), Ki67 (rabbit IgG, 1:400, Abcam), cleaved caspase-3 (rabbit IgG, 1:1000, Cell Signaling), and DCX (guinea pig IgG, 1:1000, Millipore), followed by appropriate species-specific secondary antibodies (Molecular Probes). DAPI (10 µg/ml, Sigma) was included in the secondary antibody incubations to label nuclei. The stained cells were then mounted in mounting medium and analyzed by conventional or confocal fluorescence microscopy.

For immunohistochemistry, the target region of the spinal cord (∼0.5 cm in length) was surgically dissected after perfusion, fixed in 4% paraformaldehyde (PFA) in PBS for 1 day, cryo-protected in 30% sucrose solution for 1 day, and sectioned into 25 μm horizontal slices using a Leica CM1950 cryostat. The spinal cord sections were collected serially onto Superfrost™ Plus glass slides (Fisher Scientific) and air-dried. Sections were washed in PBS three times for 5 min per wash, permeablized with 2% Triton X-100 in PBS for 20 min and blocked using a 5% normal donkey serum (NDS) and 0.1% Triton-X in PBS for 2 h to reduce non-specific binding of the antibodies. The samples were then incubated with primary antibodies diluted in the same blocking buffer at 4°C for overnight, washed in PBS three times for 5 min per wash, and incubated with secondary antibodies diluted in blocking buffer for 1 h. Finally, the samples were washed in PBS three more times for 10 min per wash and mounted with coverslips using anti-fading mounting solution (Invitrogen). DAPI (10 µg/ml, Sigma) was included in the secondary antibody incubations to label nuclei. The immunostained sections were examined and imaged using conventional or confocal fluorescence microscopes.

### Western blot analysis

Cell cultures were harvested in RIPA buffer (Alfa Aesar) following manufacturer’s instructions. Protein concentrations were determined by Coomassie Plus (Bradford) Assay Kit (Thermo Scientific). Forty µg of protein were boiled in SDS sample buffer for 5 min and loaded on each lane of Any kD™ Mini-PROTEAN® TGX™ precast polyacrylamide gels and transferred onto PVDF membranes. Western blot analysis was performed as previously described (Zhang et al., 2015). The primary antibodies were anti-ELAVL4 (mouse IgG, 1:200, Abcam), anti-ELAVL2 (rabbit IgG, 1:100, Santa Cruz), anti-ELAVL2/4 (rabbit IgG, 1:500, Abcam), and anti-GFP (rabbit IgG, 1:500, Abcam). The primary antibodies were detected by appropriate species-specific DyLight 700 or 800-conjugated secondary antibodies (1:10,000, Thermo Scientific). Anti-GAPDH (mouse IgG, 1:1000, Sigma) was used to normalize sample loadings. Quantification of relative protein expression levels was done by measuring signal intensity of target bands on a LI-COR Odyssey® Infrared Imaging System and normalizing to that of GAPDH.

### Total RNA extraction and quantitative reverse transcriptase PCR (qRT-PCR)

The procedures for total RNA extraction and qRT-PCR analysis have been previously described (Zhang et al., 2015). Briefly, total RNAs were purified from cell cultures by TRIzol® reagent following manufacturer’s instructions (Invitrogen). Quality and concentration of total RNA samples were examined on Nanodrop 8000 (Thermo Scientific). RT reactions were carried out with 0.5 µg of total RNAs using qScript^TM^ cDNA SuperMix (Quanta Biosciences). In the case of detecting miRNA expression, a Poly-A addition step was included and RT reactions were carried out with 0.5 µg of total RNAs using a miR-RT-primer (caggtccagtttttttttttttttvn) following a protocol previously described (Balcells et al., 2011). PCR was performed on a BIO-RAD CFX96™ Real-time System (BIO-RAD) using SYBR Select Master Mix (Applied Biosystems). Primer sequences for qRT-PCR are shown in Table 1. The relative gene expression levels were normalized to that of the housekeeping gene *GAPDH*. MiRNA-specific primers were designed according to a strategy previously described (Balcells et al., 2011) and shown in Table 1. The relative expression levels of individual miRNAs were normalized to that of *U6* snRNA.

**Table 1.**
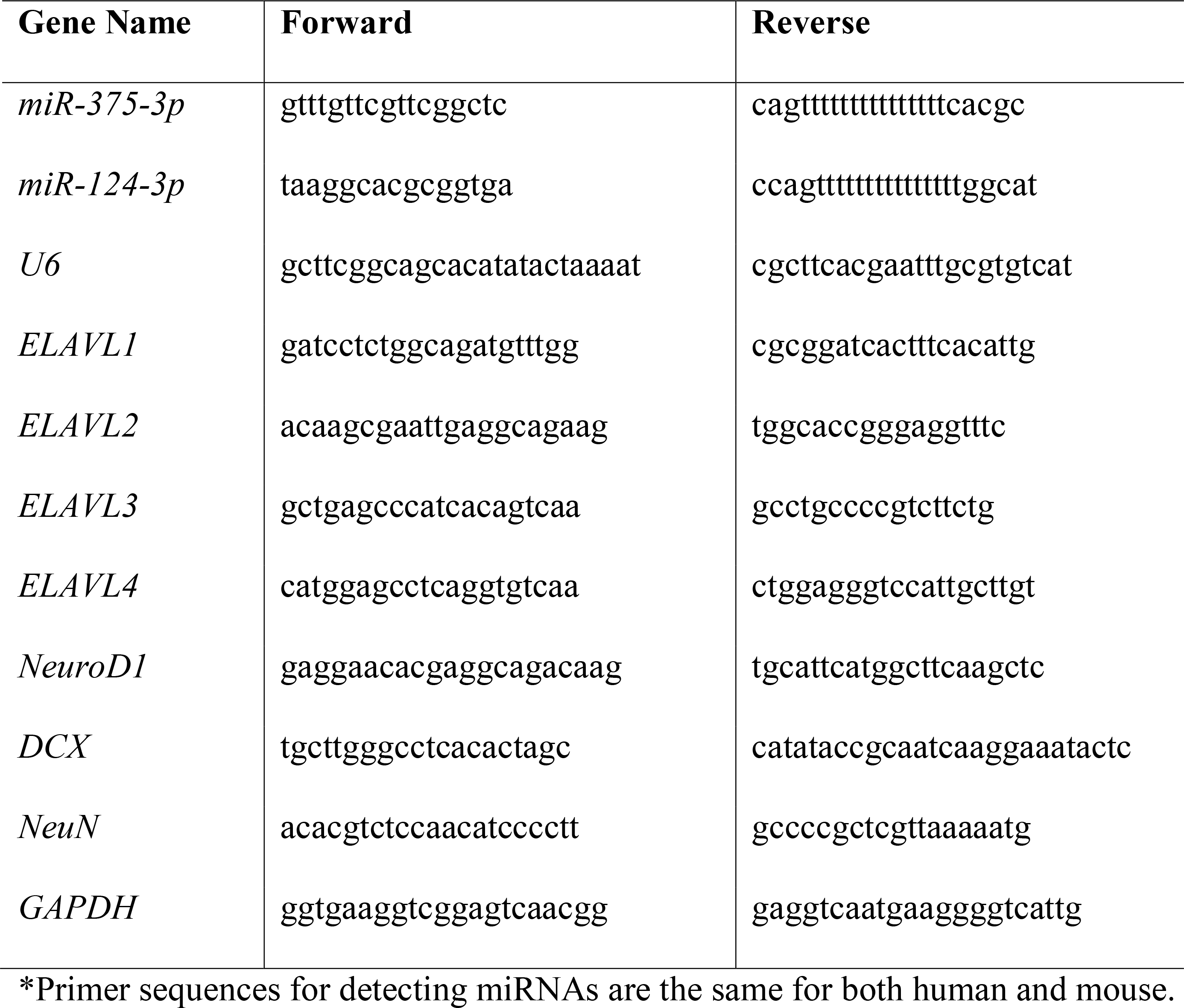
Human primers used for qRT-PCR.

### Measurements and statistical analysis

The levels of cellular fluorescence from fluorescence microscopy images were determined in ImageJ software and corrected to mean fluorescence of background readings. For quantifications on western blots, data were collected from at least three biological replicates. The data are presented as mean ± SEM. Statistical analysis was performed in GraphPad Prism 9 using Student’s *t* test for data involving only two groups, and one-way analysis of variance (ANOVA) with Bonferroni *t*-test for data involving more than two groups. Pearson’s Chi-squared test was performed in Fig. 1H. P < 0.05 was considered a significant difference.

**Figure 1.**
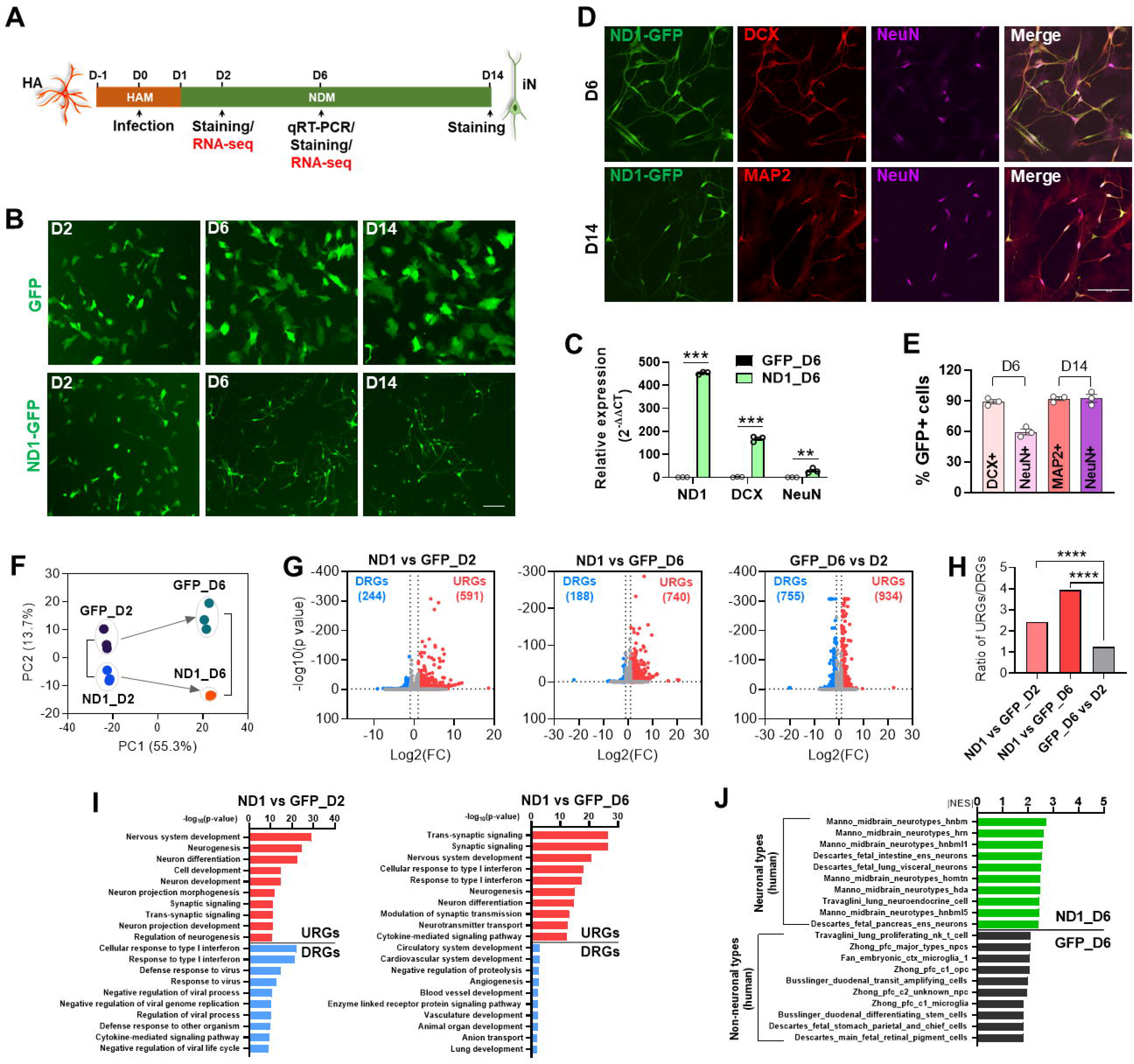
NeuroD1 activates downstream neuronal genes to initiate AtN reprogramming. (A) Schematic and timeline of NeuroD1 (ND1)-mediated neuronal reprogramming in HA culture. (B) Representative live images of morphological changes of HA transduced with GFP or ND1-GFP retroviruses at 2 (D2), 6 (D6), and 14 (D14) DPI. Scale bar, 100 μm. (C) qRT-PCR results showing expression levels of ND1, DCX, and NeuN in infected HA cultures at 6 DPI. (D) Representative staining images of ND1, DCX, MAP2, and NeuN in GFP-or ND1-GFP-infected cells in HA. Scale bar, 100 μm. (E) Quantitative analysis of (D). (F) Principal component analyses (PCA) on top 1,000 detected mRNAs from RNA-seq data. (G) Volcano plot analyses between indicated samples showing upregulated (URGs) and downregulated (DRGs) genes by ND1 as well as GFP control samples at 2 and 6 DPI. (H) Ratios of URGs and DRGs from (G). (I) Gene ontology (GO) analysis of URGs and DRGs at different time points. The top 10 GO terms in Biological Processes are displayed. (J) Gene set enrichment analysis (GSEA) of cell type signature gene sets curated from cluster markers identified in single-cell sequencing studies of human tissue. Top 10 gene sets enriched in ND1-infected cells at 6 DPI are displayed. NES, normalized enrichment score; hnbm, human medial neuroblast; hrn, human red nucleus; hnbml, human mediolateral neuroblasts; ens, enteric nervous system; homtn, human oculomotor and trochlear nucleus; hda, human dopaminergic neurons; npcs, human neural progenitor cells; pfc, prefrontal cortex; opc, oligodendrocyte progenitor cells. Data represent mean ± SEM in (C) and (E). **, p <0.01; ***, p<0.001; ****, p<0.0001.

## RESULTS

### NeuroD1 activates downstream neuronal genes to initiate AtN reprogramming

To reveal the molecular mechanisms underlying AtN reprogramming, we performed RNA-seq analysis on HA cultures during NeuroD1-mediated reprogramming. Specifically, HA cultures were infected by a NeuroD1-expressing retrovirus at day 0 followed by medium change (from HAM to NDM) to support neuronal reprogramming. Samples were harvested at different days post infection (DPI) for characterizations and analyses including RNA-seq at 2 and 6 DPI (Fig. 1A). Florescence live imaging indicated that morphological changes occur as early as 2 DPI by NeuroD1 compared with the GFP control, and that neuronal morphology becomes apparent at 6 and 14 DPI (Fig. 1B). The expression of NeuroD1 as well as the early neuronal marker DCX and the mature neuronal marker NeuN was detected during reprogramming by qRT-PCR (Fig. 1C). Furthermore, the high efficiency of NeuroD1-mediated neuronal reprogramming in HA cultures was confirmed by the high percentages of immunoreactive cells for DCX, NeuN, and MAP2 (another mature neuronal marker) at 6 and 14 DPI (Fig. 1D and 1E).

For RNA-seq, two separate libraries were prepared on each total RNA sample with one being for mRNAs to detect coding genes, and the other for small RNAs to detect miRNAs (see Experimental Procedures). Global gene expression profile undergoes drastic changes during neuronal reprogramming. As demonstrated by PCA, while replicates were clustered together, NeuroD1 and GFP control groups were already separated at 2 DPI and even more so at 6 DPI indicating a reprogramming progression over time at the global gene expression level (Fig. 1F). Differentially expressed genes (DEGs) were obtained by comparing NeuroD1 samples to GFP control samples at different time points. Interestingly, among the DEGs, the number of up-regulated genes (URGs) was much greater than that of down-regulated genes (DRGs) at both 2 and 6 DPI while a comparison between the two GFP control samples (i.e., GFP_D6 vs D2) showed less of a difference (Fig. 1G and 1H). This is consistent with the fact that NeuroD1 is a transcription activator turning on neuronal genes to initiate AtN conversion program (Ma et al., 2022). Similar patterns for the number of URGs and DRGs were also reported in ES cells and microglial cultures during NeuroD1-mediated neuronal reprogramming (Matsuda et al., 2019; Pataskar et al., 2016). In contrast, chemically induced neuronal reprogramming in the same HA cultures showed equivalent number of URGs and DRGs, at least during first week of induction (Ma et al., 2019). These results further demonstrate the instructive role of NeuroD1 as a reprogramming factor.

We also performed Gene Ontology (GO) analysis to identify major functional categories of the DEGs. As expected, the GO terms of URGs are mostly neuron-related ones containing “neurogenesis”, “development”, and “synaptic” (Fig. 1I). Interestingly, we observed more terms containing “development” in URGs at D2 and more terms containing “synaptic” at D6 suggesting a progression from immature to mature neuronal state during reprogramming (Fig. 1I). In contrast, the GO terms of DRGs are all non-neuronal ones including interferon and virus-related ones, which could be due to differential cellular responses to retrovirus infection between different conditions (Fig. 1I). We then performed Gene Set Enrichment Analysis (GSEA) of cell type signature gene sets with single-cell sequencing data of human tissue. The results showed that top 10 gene sets enriched in the NeuroD1_D6 sample are various types of human neurons, while top 10 gene sets enriched in the GFP_D6 sample are all non-neurons (Fig. 1J). This further confirms neuronal reprogramming by NeuroD1 in HA cultures.

### Identification of NeuroD1-induced “core” genes during glia-to-neuron reprogramming

NeuroD1 has been utilized to reprogram other glial cell types including microglia to neurons, and drive ES cells into neurons when overexpressed (Matsuda et al., 2019; Pataskar et al., 2016). To identify conserved genetic pathways during NeuroD1-mediated neuronal reprogramming, we compared our RNA-seq data with publicly available datasets from others and found an overlap of 70 genes, all of which are URGs, across 4 datasets (Fig. 2A). There were no overlapping genes among the DRGs likely due to the different gene expression profiles of the original cell types. This overlapping pattern among URGs and DRGs between different cell types further indicates that NeuroD1 induces neuronal differentiation and/or reprogramming mainly through activating consensus genetic programs. Functional annotation clustering analysis showed that these overlapping “core” genes possess unique GO terms contributing to the neuronal reprogramming process (Fig. 2B). These include “synapse” and “glutamatergic synapse”, which are essential functional structures of mature neurons and are consistent with the fact that NeuroD1-reprorammed neurons are mostly glutamatergic subtypes (Chen and Li, 2022; Guo et al., 2014). Other GO terms such as “neurogenesis” and “neuron projection morphogenesis” indicate the developmental process of neuronal differentiation. “Axons” and “plasma membrane” contain important genes such as potassium-channels *KCNC1*, *KCNQ2*, and sodium channel *SCN3A*, that are critical to functional properties of neurons. There are also a group of genes encoding “RNA binding” proteins, suggesting the importance of post-transcriptional regulation during neuronal reprogramming process (Fig. 2B). Gene network analysis of the core genes by STRING program showed that most of these genes are interconnected, with top representations of GO terms being “neurogenesis”, “synapse”, “excitatory synapse”, and “RNA recognition motif” (Fig. 2C). Identification of the core genes will help define a molecular signature of NeuroD1 action during neuronal reprogramming.

**Figure 2.**
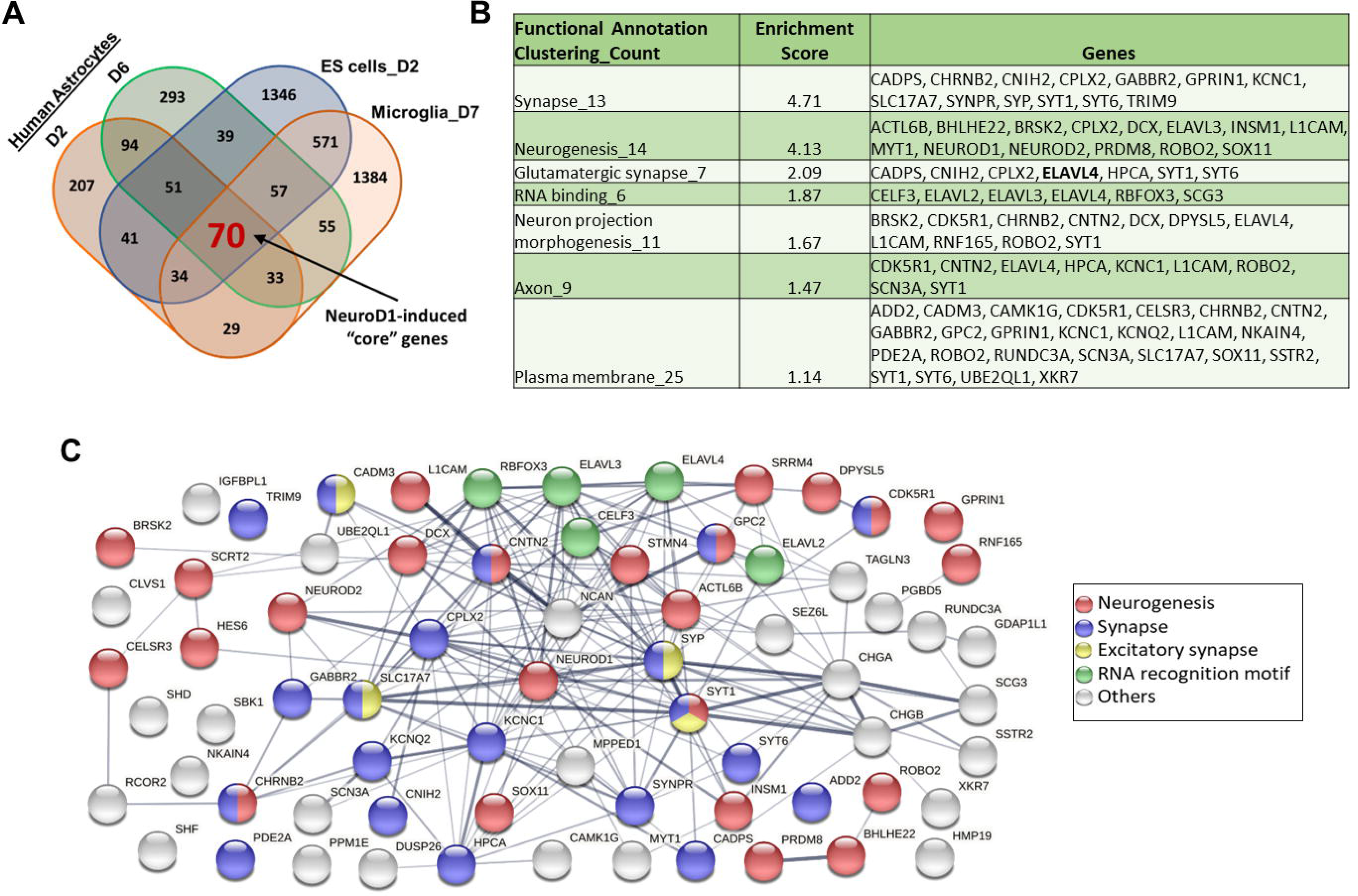
Identification of 70 core genes induced by NeuroD1 during neuronal reprogramming and differentiation. (A) Venn diagram showing the overlap between our gene sets and published gene sets that are upregulated during ND1 induced embryonic stem (ES) cells-to-neuron differentiation (Pataskar A, et al. EMBO J. 2016) and microglia-to-neuron reprogramming (Matsuda T, et al. Neuron. 2019). (B) Table showing the selective functional annotation clusters of 70 overlapping genes by DAVID. (C) Most of the proteins encoded by 70 core genes are interconnected based on the STRING database.

### NeuroD1 induces miRNA expression changes during AtN reprogramming

To determine miRNA function during AtN reprogramming, we first examined expression changes of miRNAs in the small RNAs portion of our RNA-seq data. In PCA, miRNA profiles of NeuroD1 and GFP control groups are clustered closely at D2 and diverged at D6 (Fig. 3A), like the profiles of mRNAs (Fig. 1F). A small number of differentially expressed miRNAs (DEmiRs) were identified by comparing NeuroD1 and GFP groups at both time points as well as by comparing the two GFP groups. Like mRNAs, there are more upregulated miRNAs (URmiRs) (5 at D2 and 19 at D6) than downregulated miRNAs (DRmiRs) (1 at D2 and 9 at D6) by NeuroD1 at both time points as shown by volcano plots (Fig. 3B). The two most significant URmiRs during NeuroD1-mediated neuronal conversion in HA based on *p*-value and fold change are hsa-miR-375-3p (miR-375) and hsa-miR-124-3p (miR-124) (labeled red in Fig. 3C). Based on average read counts (RC), the highly expressed miRNAs in HA cultures include miR-21-5p that has been shown to function in astrocytes (Bhalala et al., 2012; Liu et al., 2018) (Fig. 3D). However, these highly expressed miRNAs are not changing their expression during AtN reprogramming. In contrast, both miR-375 and miR-124 have a minimal expression level in HA and are induced drastically by NeuroD1 (Fig. 3D) suggesting their important functions during the reprogramming process. We further confirmed the expression changes of these two miRNAs by qRT-PCR (Fig. 3E). Of note, the detection of significant upregulation of miR-124 at D2 by qRT-PCR, but not by RNA-seq, indicates a higher sensitivity of qRT-PCR method.

**Figure 3.**
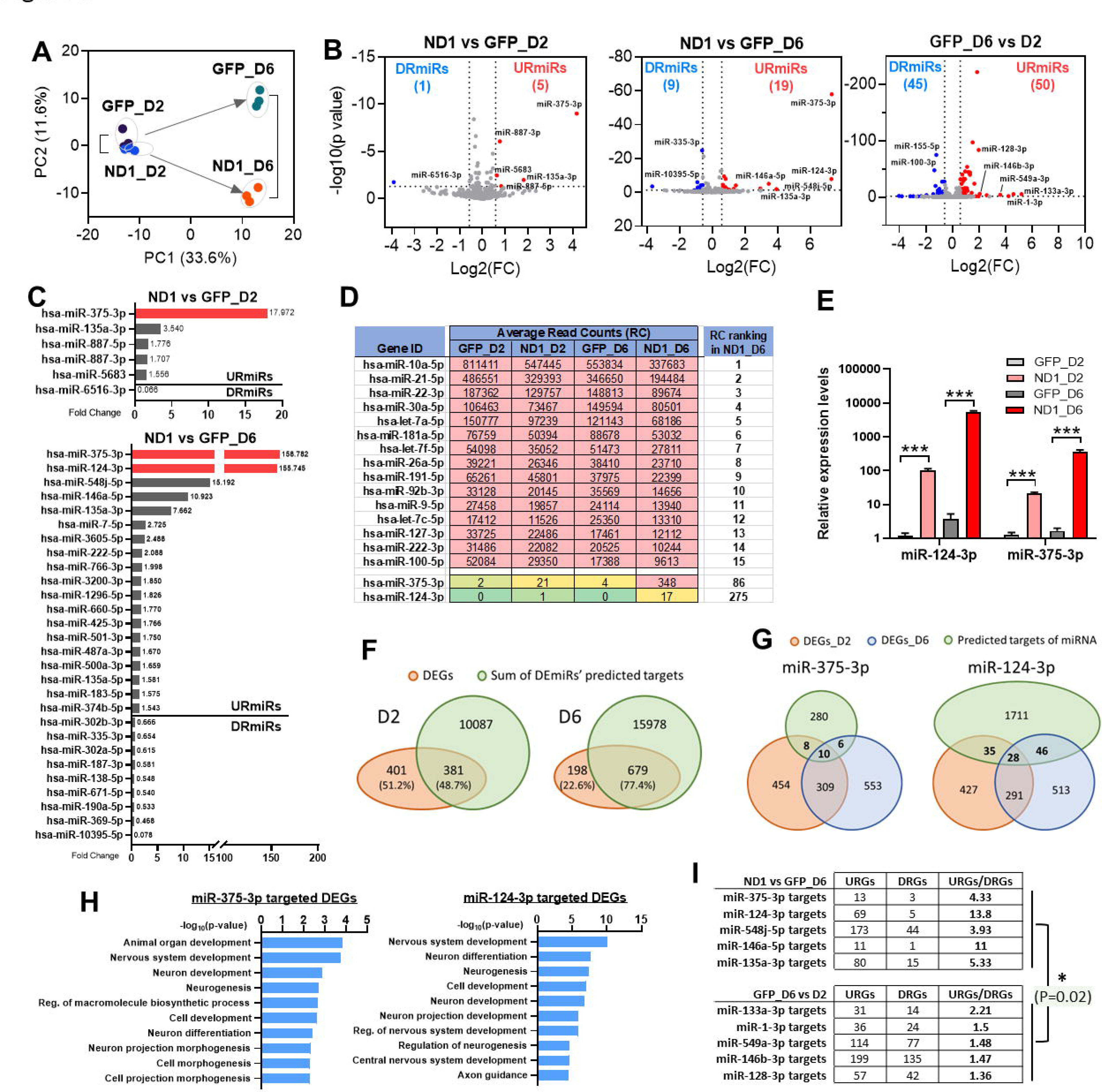
NeuroD1 induces expression changes of miRNAs that are predicted to target DEGs during AtN reprogramming. (A) Principal component analysis (PCA) on all detected miRNAs in HA. (B) Volcano plots of the upregulated (URmiRs) and downregulated (DRmiRs) miRNAs during ND1-mediated AtN reprogramming at 2 and 6 DPI. (C) Fold change ranking of URmiRs and DRmiRs in ND1-infected HA at 2 and 6 DPI as compared with GFP controls. The two most upregulated miRNAs, has-miR-375-3p and has-miR-124-3p, are highlighted in red. (D) Table showing top 15 highly expressed miRNAs in “ND1_D6” plus hsa-miR-375-3p and hsa-miR-124-3p with average read counts (RC) and ranking. (E) qRT-PCR analysis of miR-124-3p and miR-375-3p expression during AtN reprogramming process. (F) Venn diagrams showing the overlap of DEGs and predicted targets of all DEmiRs downloaded from TargetScan database (v7.2). (G) Venn diagrams showing the overlap of DEGs and predicted targets of miR-375-3p and miR-124-3p downloaded from TargetScan database (v7.2). (H) Top 10 GO terms of miR-375-3p and miR-124-3p targeted DEGs. (I) Comparison of the number and ratio of miRNA targeted URGs and DRGs between “ND1 vs GFP_D6” and “GFP_D6 vs D2” conditions. Data represent mean ± SEM in (E). *, p<0.05; ***, p<0.001.

To determine the impact of DEmiRs on the global gene expression changes, we extracted predicted target genes of these DEmiRs by *TargetScan* program and overlaid them with DEGs. About half of DEGs (381/782) are predicted DEmiR targets at D2, and even more (679/877) at D6 (Fig. 3F). Therefore, DEmiRs likely make a significant contribution to the changing genetic program during AtN conversion. MiR-375 and miR-124 also show overlapping target genes with DEGs at both time points (Fig. 3G); these target genes have highly enriched neuronal GO terms (Fig. 3H). Given the inhibitory mechanism of action for miRNAs on gene expression, one would expect an inverse correlation of expression levels between DEmiRs and their predicted DEG targets. Surprisingly, most of the target DEGs of URmiRs are URGs during AtN reprogramming, and the ratios of target URGs/DRGs from ND1 vs GFP_D6 are significantly higher than those from GFP_D6 vs D2 (Fig. 3I). These results suggest that when overexpressing NeuroD1 as a strong transcriptional driver for gene expression during AtN reprogramming, one major function of DEmiRs could be to modulate DEGs expression to ensure an optimal level for successful reprogramming.

### MiR-375 regulates *nELAVLs* expression level during NeuroD1-mediated AtN reprogramming

While there is an abundance of literature on the role of miR-124 in neuronal differentiation and reprograming (Cheng et al., 2009; Lu et al., 2021; Papagiannakopoulos and Kosik, 2009; Sanuki et al., 2011; Wohl et al., 2019), relatively little is known about miR-375’s contribution to the same processes. Therefore, we decided to investigate the function of miR-375 during NeuroD1-mediated neuronal reprogramming in our HA culture system. There are only 10 miR-375 targets overlapping with DEGs between two time points (Fig. 3G); these targets are listed with their mean expression levels from different samples by RNA-seq (Fig. 4A). Among the very few miR-375 targeted DEGs, we identified three members of an RNA-binding protein family (*ELAVL*), i.e. *ELAVL2, 3*, and *4* (Fig. 4A). While *ELAVL1* is regarded as universally expressed by many cell types, *ELAVL2-4* are preferentially expressed in neurons (Bronicki and Jasmin, 2013). Interestingly, the 3’-UTRs of *ELAVL2-4*, but not *ELAVL1*, contain one or more predicted miR-375 binding sites potentially allowing gene expression regulation by miR-375 (Fig. 4B). Indeed, *ELAVL4* has been experimentally confirmed as miR-375 target in other systems (Abdelmohsen et al., 2010; Samaraweera et al., 2014). We first confirmed *ELAVL2-4* gene expression changes by qRT-PCR (Fig. 4C) and changes of ELAVL2 and 4 at protein level by western blot (Fig. 4D) during AtN reprogramming. ELAVLs as RNA-binding proteins stabilize target mRNAs for longer half-life and thereby increase protein translation (Anderson et al., 2001). ELAVLs recognize an AU-rich element (ARE) sequence allowing prediction of their target genes (Bronicki and Jasmin, 2013). Since three *ELAVLs* are highly upregulated during NeuroD1-mediated reprogramming, we wanted to see how many of the ARE genes are also upregulated that are potentially the result of elevated *ELAVLs*. Indeed, more than 26% of the DEGs are predicted ARE genes at both time points, and their GO terms are enriched in neuronal ones (Fig. 4E).

**Figure 4.**
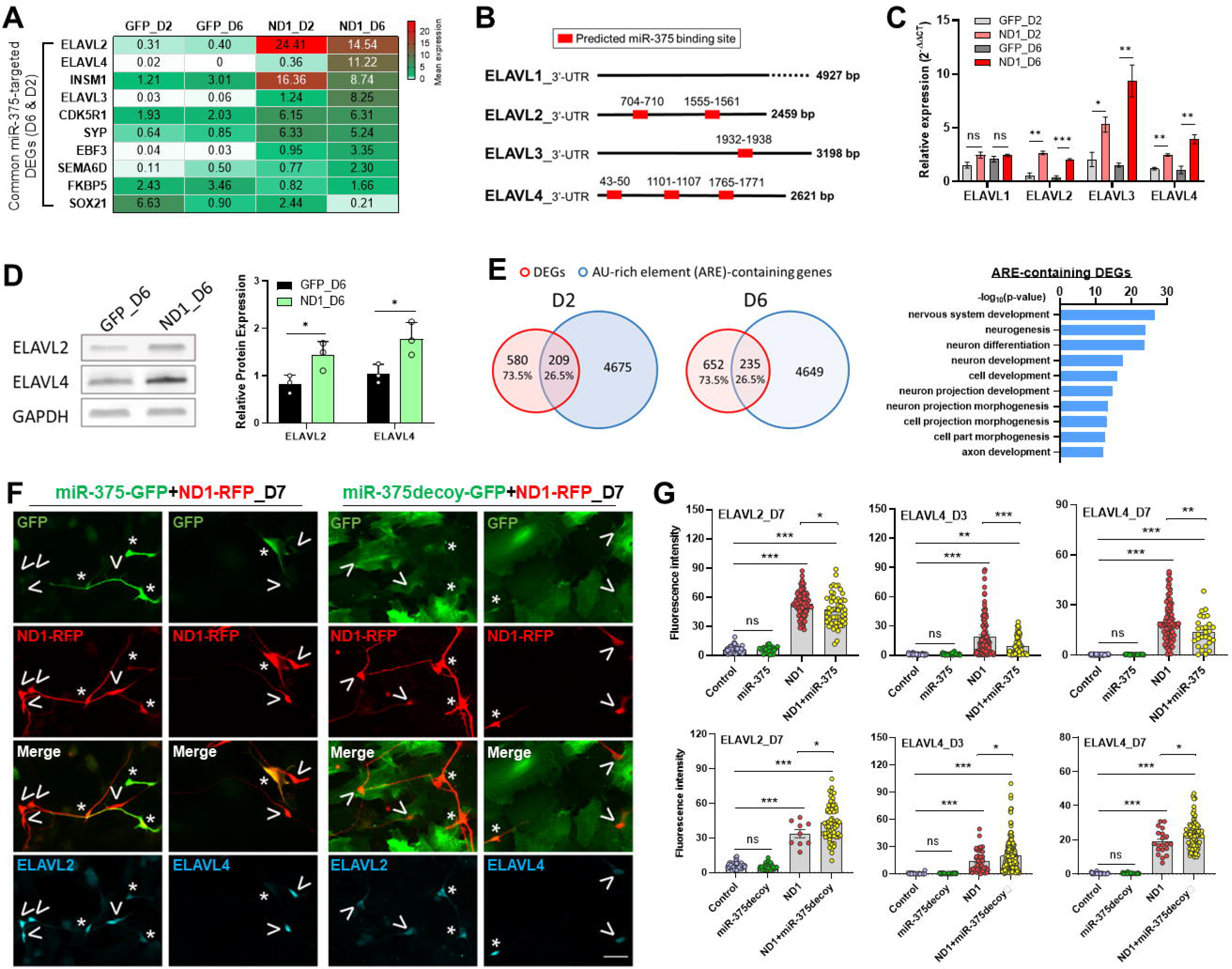
MiR-375 regulates protein expression level of nELAVLs during NeuroD1-mediated AtN reprogramming. (A) Heatmap representation of the 10 common predicted targets of miR-375-3p overlapped with DEGs at both 2 and 6 DPI. Normalized expression levels are also presented. (B) Computational prediction of miR-375-3p target sites in the 3’-UTRs of ELAVLs from TargetScan. (C) qRT-PCR analysis of *ELAVLs* mRNA expression during <u>AtN</u> reprogramming in HA. (D) Immunoblot analysis and quantification of ELAVL2 and ELAVL4 protein in GFP-or ND1-infected HA cultures at 6 DPI. (E) Venn diagrams of DEGs overlapped with AU-rich element (ARE) genes and the top 10 GO terms of ARE-containing DEGs. (F) Representative immunostaining images of GFP, RFP, ELAVL2 and ELAVL4 in HA coinfected with miR-375-GFP + ND1-RFP and miR-375decoy-GFP + ND1-RFP viral constructs. >, ND1 alone-infected HA; *, coinfected HA. Scale bar, 20 μm. (G) Quantitation of ELAVL2 and ELAVL4 protein immunofluorescence intensity of coinfected HA in (F). Data represent mean ± SEM. *, p<0.05; **, p<0.01; ***, p<0.001; ns, not significant.

To determine if miR-375 can regulate *nELAVLs* gene expression level during NeuroD1-mediated reprogramming, we adopted overexpression and knockdown strategies. For overexpression, we constructed a retrovirus expressing miR-375 under the CAG promoter. For knockdown, we introduced a lentivirus expressing a miR-375 inhibitor (miR-375-decoy) that has been validated to knock down miR-375 level by more than 50% (Mullokandov et al., 2012). Both viral constructs have been confirmed to be able to regulate the expression level of mature miR-375 upon infection (data not shown). Our design was to coinfect HA with each of these miR-375 constructs and NeuroD1, and compare with NeuroD1 alone to see if changing miR-375 level in the presence of NeuroD1 could induce changes in the expression level of *nELAVLs*. For this, we decided to detect changes at protein level by immunostaining. We found that, upon coinfection, overexpression of miR-375 decreases ELAVL2 and 4 protein levels in coinfected cells compared with cells infected by NeuroD1 alone, while reduction of miR-375 by decoy increases (Fig. 4F and 4G). Therefore, our data indicate that NeuroD1-induced increase of nELAVLs can be further regulated by miR-375 level; miR-375 can indirectly modulate expression of ARE genes via ELAVLs as well as directly regulate its target genes.

### MiR-375/*nELAVLs* are also induced by neuronal reprogramming factors ASCL1 and Neurog2

To determine if miR-375/*nELAVLs* upregulation is unique to NeuroD1-mediated neuronal reprogramming, we analyzed their expression in AtN induced by two other reprogramming factors ASCL1 and Neurog2. At 6 DPI, the neuronal reprogramming by three factors was confirmed by a morphological change of infected cells in HA albeit ASCL1 was less potent in reprogramming than NeuroD1 and Neurog2 (Fig. 5A and 5B). We further confirmed overexpression of these reprogramming factors in infected HA cultures by qRT-PCR (Fig. 5C). Consistent with the morphological change upon reprogramming, qRT-PCR analysis also showed induction of neuronal markers DCX and NeuN as well as *nELAVLs* by all three factors at 6 DPI except that NeuN is not significantly upregulated by ASCL1 (Fig. 5D). Furthermore, both miR-375 and miR-124 are markedly upregulated by all three reprogramming factors, interestingly with a much higher level of miR-375 induced by ASCL1 than the other two factors (Fig. 5E).

**Figure 5.**
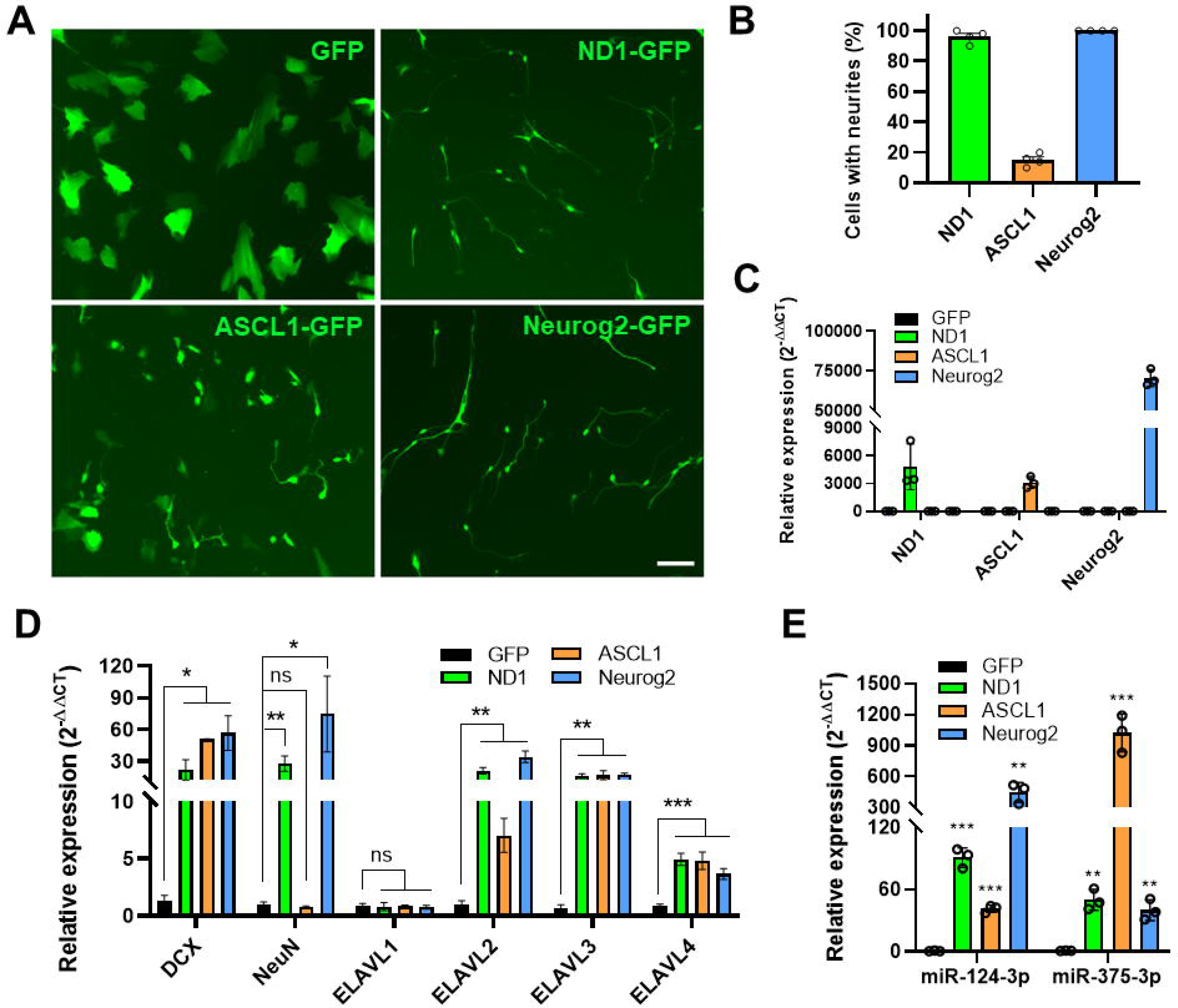
MiR-375/*nELAVLs* are induced by neuronal reprogramming factors ASCL1 and Neurog2 in HA. (A) Representative live images of morphological changes in HA cultures transduced with GFP, ND1-GFP, ASCL1-GFP, and Neurog2-GFP retroviruses at 6 DPI. Scale bar, 50 μm. (B) Quantification showing the percentage of neuron-like cells in infected HA cultures at 6 DPI. QRT-PCR analyses showing expression of the reprogramming factors (C), neuronal markers and *ELAVLs* (D), and miRNAs (E) in infected HA cultures at 6 DPI. Data represent mean ± SEM. *, p<0.05; **, p<0.01; ***, p<0.001; ns, not significant.

These data suggest that miR-375/*nELAVLs* interaction could be a common mechanism during neuronal reprogramming induced by different reprogramming factors.

### NeuroD1 induces ELAVL4 expression in the injured mouse spinal cord and miR-375/ELAVL4 expression in mouse astrocyte cultures during AtN reprogramming

We next wanted to determine if miR-375/*nELAVLs* interaction is relevant during AtN reprogramming *in vivo*. To target astrocytes in the mouse spinal cord, we applied AAV-Cre-Flex expression system as previously described (Puls et al., 2020). A mixture of AAV5-GFAP-Cre and AAV5-Flex-NeuroD1-GFP was injected into the mouse spinal cord after a stab injury. Based on our previous characterization of AtN reprogramming in the injured spinal cord, 2 week post injection (WPI) represents an intermediate state of reprogramming when we detected mostly “transitional cells” that are positive for both GFAP and NeuN (Puls et al., 2020). At 2 WPI, we first observed many AAV-infected astrocytes that are GFAP+ around the injury sites (asterisks in Fig. 6A); interestingly, AAV-ND1-GFP-infected astrocytes show smaller cell bodies than AAV-GFP-infected ones as revealed by GFP fluorescence (Fig. 6A) indicating ongoing AtN reprogramming. We also confirmed NeuroD1 expression in AAV-ND1-GFP-infected astrocytes by immunostaining (Fig. 6B) and the presence of “transitional cells” in the AAV-ND1-GFP-infected spinal cord as expected (Fig. 6C). Importantly, among the AAV-ND1-GFP-infected astrocytes, we were able to identify some cells also expressing ELAVL4 protein (arrowheads, Fig. 6D). The expression level of ELAVL4 in these cells is much weaker than the nearby neurons (Fig. 6D) suggesting an early onset of neuronal gene expression at this stage and yet becomes much stronger at later stages (i.e., 4 WPI) when the cells acquire neuronal morphology (arrowheads, Fig. 6E). To detect miR-375 expression by NeuroD1, we turned to mouse neonatal astrocyte cultures for the convenience of analysis. NeuroD1-expressing retrovirus can readily reprogram mouse cultured astrocytes into DCX+/NeuN+ neurons within 2 weeks (data not shown). We confirmed the morphological change and NeuroD1 expression of ND1-GFP-infected cells compared with GFP control at 9 DPI, and induction of ELAVL4 by NeuroD1 by immunostaining (arrows, Fig. 6F). Lastly, qRT-PCR data showed that both miR-375-3p and miR-124-3p are significantly upregulated by NeuroD1 in these astrocyte cultures (Fig. 6G). Therefore, NeuroD1 can induce miR-375/ELAVL4 expression in mouse astrocyte cultures as in HA and induce at least ELAVL4 protein in astrocytes of the injured spinal cord during AtN reprogramming.

**Figure 6.**
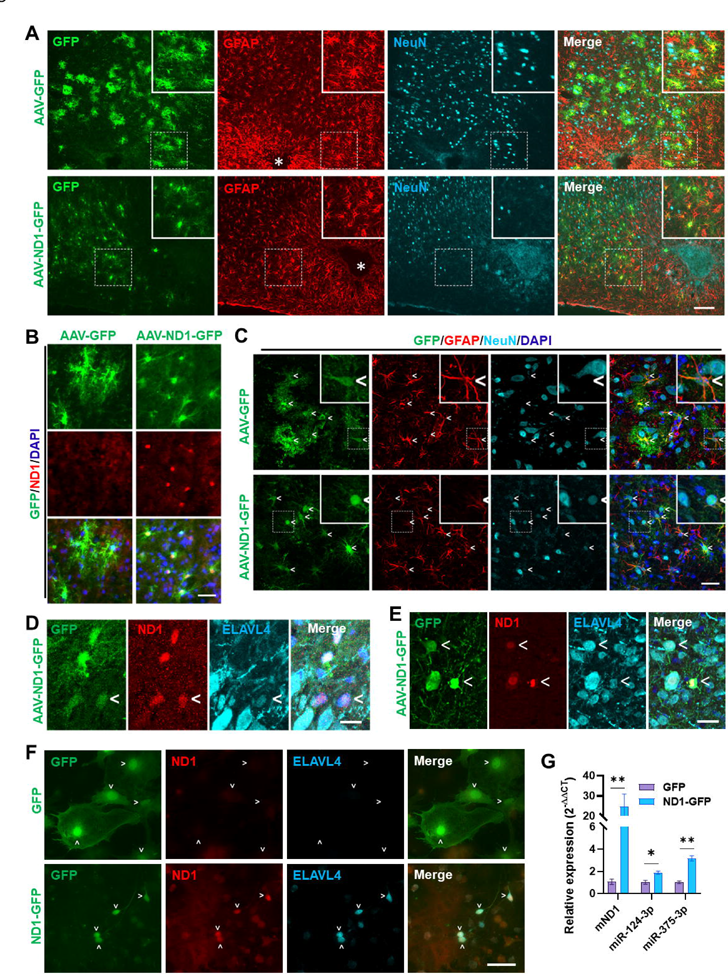
NeuroD1 induces miR-375/ELAVL4 expression in mouse astrocyte cultures and ELAVL4 expression in the injured spinal cord during AtN reprogramming. (A) Representative images of GFP, GFAP, and NeuN immunostaining in the injured mouse spinal cord at 2 weeks post injection (WPI) of a mixture of AAV5-GFAP-Cre with either AAV5-Flex-GFP or AAV5-Flex-ND1-GFP at a 1:9 ratio. *, stab injury sites. Scar bar, 100 μm. (B) Representative images of GFP and NeuroD1 immunostaining of the same experiment in (A). Scar bar, 20 μm. (C) Confocal images of (A) to show “transitional cells” that are GFAP+/NeuN+ infected by AAV5-ND1-GFP. <, AAV-infected cells. Scar bar, 20 μm. (D) Confocal images of GFP, ND1, and ELAVL4 immunostaining of the same experiment in (A). <, a ND1-GFP-infected astrocyte expresses ND1 and ELAVL4 protein. Scar bar, 10 μm. (E) Confocal images of GFP, ND1, and ELAVL4 immunostaining in the injured mouse spinal cord at 4 WPI of a mixture of AAV5-GFAP-Cre and AAV5-Flex-ND1-GFP. <, ND1-GFP-infected cells express ND1 and ELAVL4 protein. Scar bar, 10 μm. (F) Representative images of GFP, ND1, and ELAVL4 immunostaining in mouse astrocyte cultures infected with GFP or ND1-GFP retrovirus at 9 DPI. >, infected cells. Scar bar, 50 μm. (G) QRT-PCR analysis of mouse astrocyte cultures infected with GFP or ND1-GFP retrovirus at 9 DPI. The graph is presented as mean ± SEM. *, p<0.05; **, p<0.01.

### Overexpression of miR-375 elicits a protective effect during NeuroD1-mediated AtN reprogramming by reducing apoptosis

Given that miR-375 can potentially regulate many DEGs during NeuroD1-mediated reprogramming either directly or indirectly through *nELAVLs* (Figs. 3G and 4E), we set out to determine if manipulating miR-375 level would affect the AtN reprogramming outcome in HA. We set up a coinfection experiment combining NeuroD1-RFP with GFP control, miR-375-GFP or miR-375-decoy (Fig. 7A). The typical coinfection efficiency is around 50% in retrovirus combinations and much higher with lentivirus (i.e., miR-375-decoy) (Fig. 7B). We realized that many miR-375-GFP retrovirus-infected cells exhibit a much weaker GFP signal than control GFP virus-infected ones (Fig. 7A). The GFP signal of miR-375-GFP becomes so weak in long-term cultures and even cannot be reliably detected by immunostaining. Therefore, we performed our quantitative analyses among NeuroD1-infected cells in all the coinfection experiments without distinguishing singly and doubly infected cells. We examined AtN reprogramming by immunostaining (Fig. 7C) and showed that reprogramming efficiency as measured by the neuronal markers DCX and NeuN did not differ among the coinfected cultures (Fig. 7D), and that 2-fold more marker-positive cells per field were observed in miR-375 coinfection than GFP control (Fig. 7E). The knockdown of miR-375 by the decoy construct in the presence of NeuroD1 did not show a significant difference from the control coinfection by the analyses mentioned above (Fig. 7E). Live imaging also indicated that there were significantly more NeuroD1-infected cells in the miR-375 coinfected HA culture than in other coinfections at 28 DPI, even though all cultures started with equivalent numbers of infected cells at 2 DPI (Fig. 7F and 7G). QRT-PCR analysis showed that miR-375-GFP retrovirus induces a much higher level of miR-375 than NeuroD1+GFP at both early and late stages (Fig. 7H). Therefore, simultaneous miR-375 overexpression resulted in more NeuroD1-converted neurons in HA cultures.

**Figure 7.**
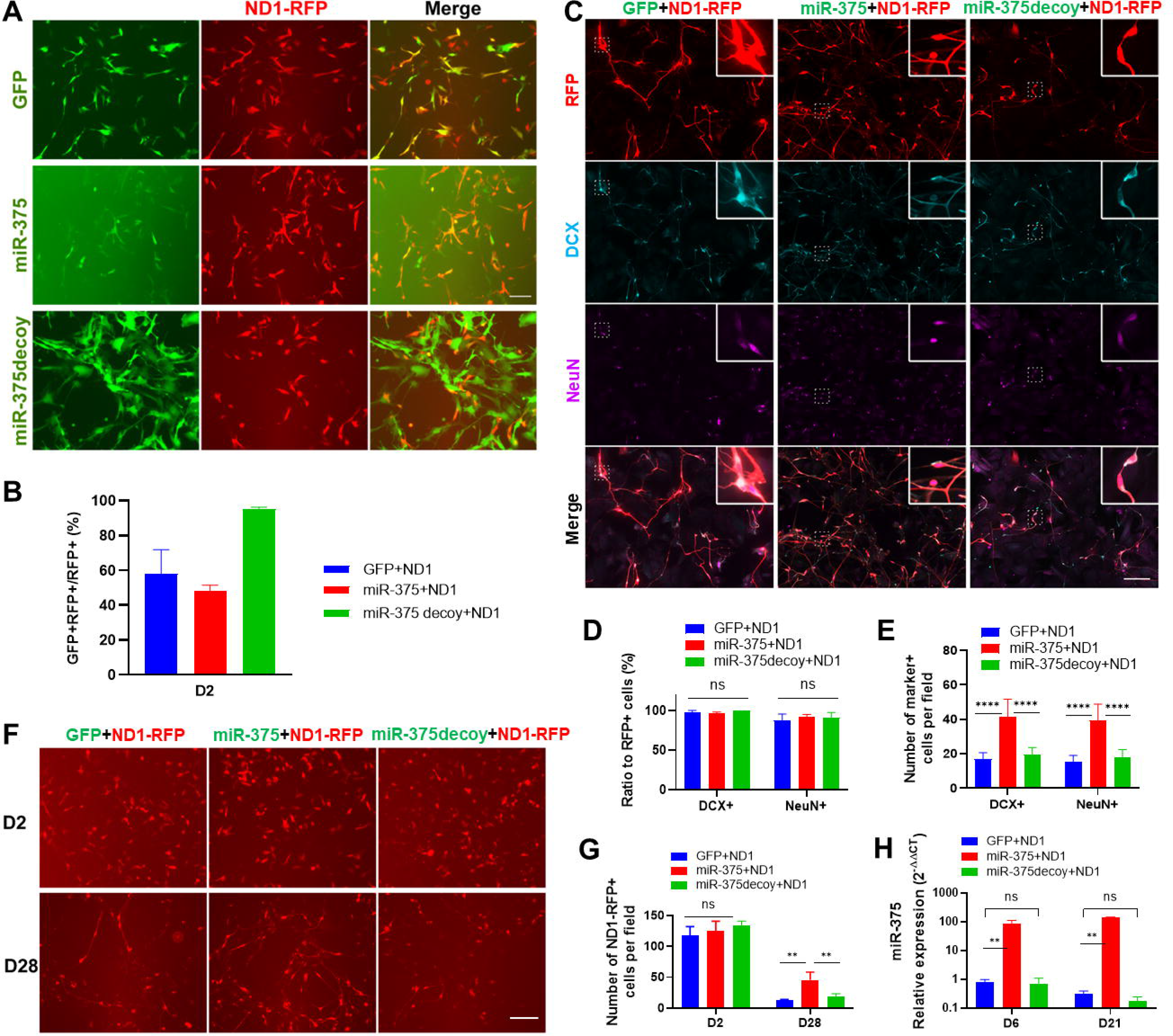
Overexpression of miR-375 increases the number of NeuroD1-reprogrammed neurons in HA. (A) Representative live images of HA coinfected by ND1-RFP with GFP, miR-375-GFP or miR-375decoy-GFP viruses at 2 DPI. Scale bar, 100 μm. (B) Quantification showing the percentage of coinfected HA with both GFP and RFP signals at 2 DPI (D2). (C) Representative immunostaining images of RFP showing ND1-RFP-infected HA, DCX, and NeuN in coinfection HA cultures at 14 DPI. Scale bar, 100 μm. Quantitative analysis of the conversion efficiency (D) and the number of converted neurons with DCX+ or NeuN+ (E) in (C). (F) Representative RFP live images of coinfected HA cultures as indicated at 2 DPI (D2) and 28 DPI (D28). Scale bar, 100 μm. (G) Quantification showing the number of ND1-infected cells (RFP+) per field at different time-points in (F). (H) qRT-PCR analysis of miR-375-3p expression in coinfected HA at early and late time points. Data represent mean ± SEM. **, p<0.01; ****, p<0.0001; ns, not significant.

We reasoned that the increased number of converted neurons in the miR-375 coinfected HA cultures could be due to decreased apoptosis and/or increased cell proliferation during reprogramming process. We first assessed cell proliferation during reprogramming in HA cultures with anti-Ki67 antibody. As expected, NeuroD1 as a neuronal differentiation factor inhibited cell proliferation and drastically decreased Ki67+ cell number at 3, 7 and 14 DPI when compared with uninfected cells, and yet no difference was observed between the two coinfection cultures (Fig. 8A and 8B). To get an overview of apoptosis status during NeuroD1-mediated neuronal reprogramming, we stained the NeuroD1-infected HA cultures with an antibody against cleaved caspase 3 (CCasp3). We found that ∼8% of NeuroD1-infected cells were CCasp3+ at 3 DPI and this percentage dropped down to a minimal level at 7 and 14 DPI (Fig. 8C and 8D). Surprisingly, miR-375 coinfected HA cultures showed a drastic decrease in the number of CCasp3+ cells when compared with GFP coinfected ones at 3 DPI (Fig. 8D), suggesting a cell protective effect of miR-375 at early stages of reprogramming. To mimic the stressed condition in disease/injury, we exposed the reprogramming HA culture to the chemotherapeutic agent Cisplatin (Yu et al., 2018). First, we treated normal HA with various doses of Cisplatin and showed a dose-response of HA in expressing CCasp3 (data not shown). We then treated NeuroD1-infected HA cultures at 2 DPI with a low and high dose of Cisplatin (5 and 250 μM, respectively) for 24 h. Strikingly, at both doses of Cisplatin treatment, overexpression of miR-375 reduced not only the percentage of CCasp3+ cells (Fig. 8E and 8F) but also the cellular CCasp3 expression level as measured by florescence intensity when compared with control groups (Fig. 8G).

**Figure 8.**
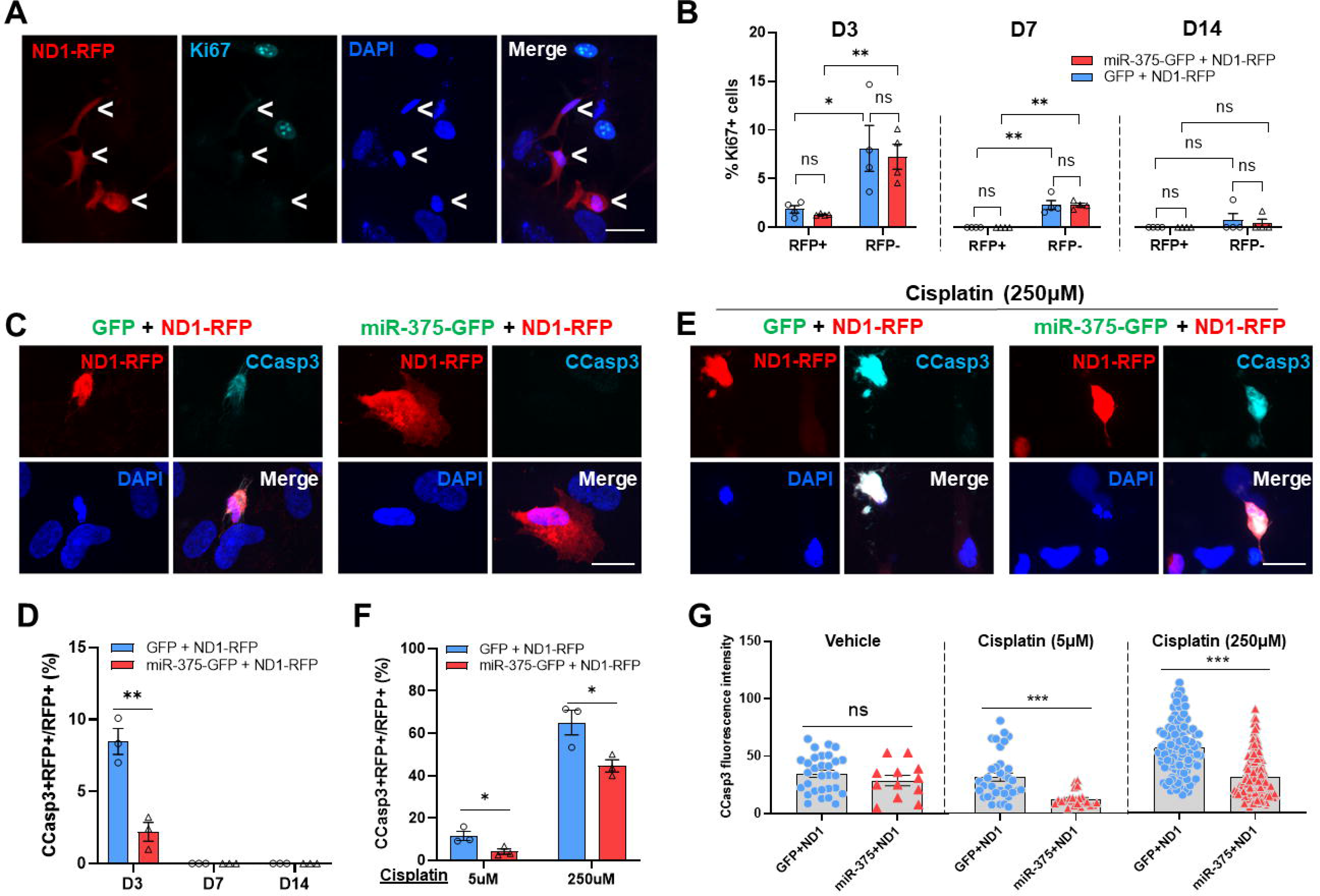
Overexpression of miR-375 reduces apoptosis during AtN reprogramming. (A) Representative immunostaining images of Ki67 at 3 DPI. <, ND1-RFP-infected cells. Scale bar, 25 μm. (B) Graphs showing the proportion of Ki67+ HA in the ND1-infected and uninfected cells at different time points post infection. (C) Representative immunostaining images of cleaved caspase-3 (CCasp3) in GFP+ND1-or miR-375+ND1-coinfected HA at 3 DPI. Scale bar, 25 μm. (D) Quantification showing the proportion of CCasp3+/RFP+ cells in the ND1-infected HA (RFP+) at different time points post infection. (E) Representative immunostaining images of CCasp3 in infected HA at 3 DPI treated with 250 μM of Cisplatin for 24 h. Scale bar, 25 μm. (F) The proportion of CCasp3+/RFP+ cells in ND1-infected HA (RFP+) at 3 DPI treated with 5 μM and 250 μM of Cisplatin for 24 h. (G) Quantitation of CCasp3 immunofluorescence intensity of CCasp3+ infected HA treated with vehicle, 5 μM and 250 μM of Cisplatin. Data represent mean ± SEM. *, p<0.05; **, p<0.01; ****, p<0.0001; ns, not significant.

To assess the potential long-term effect of miR-375 overexpression in reprogrammed neurons, we cultured these neurons for 30 days. Since the miR-375 overexpressing retrovirus loses reporter GFP expression over time, we enriched double-infected HA by FACS at 3 DPI when the miR-375-GFP level is still high (Fig. 9A). The sorting efficiency is quite high, and we detected almost 100% coinfection efficiency in the ND1-RFP+/miR-375-GFP+ group one day after plating while no GFP signal was observed in the ND1-RFP+ group (Fig. 9B). After 30 days in culture, we examined the expression of ELAVL4 and the mature neuronal markers MAP2, SYP, and NeuN in these reprogrammed neurons by immunostaining (Fig. 9C). We found that the levels of these markers are not reduced but even increased in ND1+miR-375-reprogrammed neurons compared with ND1-reprogrammed ones (Fig. 9D), suggesting that miR-375 overexpression doesn’t interfere with neuronal maturation. The slight but significant increase of ELAVL4, MAP2, and SYP levels is likely due to healthiness of neurons in the double-infected group. Taken together, miR-375 overexpression improves NeuroD1-mediated AtN reprogramming efficiency in HA cultures by inhibiting apoptosis at early stages and doesn’t compromise maturation of reprogrammed neurons in long term cultures.

**Figure 9.**
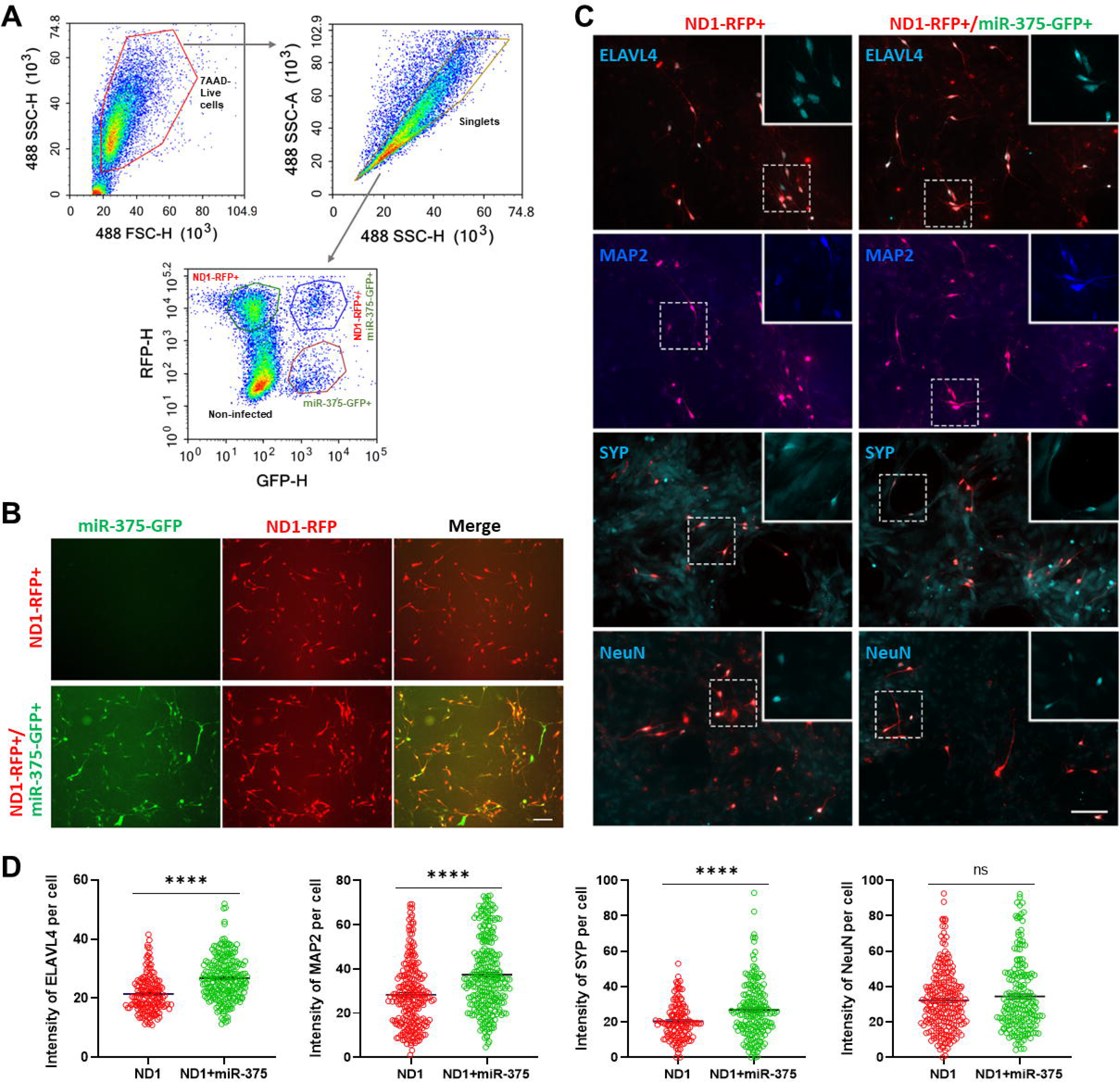
Overexpression of miR-375 does not inhibit expression levels of ELAVL4 and other mature neuronal markers of reprogrammed neurons in long-term HA cultures. (A) FACS gating strategy on dissociated HA cultures co-infected with ND1-RFP and miR-375-GFP retroviruses at 3 DPI. (B) Representative live florescence images of sorted cells that are plated 1 day after FACS. Scale bar, 100 μm. (C) Representative immunostaining images of ELAVL4, MAP2, synaptophysin (SYP), and NeuN in long-term cultures of FACS-sorted cells at 30 DPI. Scale bar, 100 μm. (D) Quantitation of immunofluorescence intensity in FACS-sorted cells in (C). ****, p<0.0001; ns, not significant.

### Overexpression of *ELAVL4* triggers apoptosis in HA and reverses miR-375-induced survival-promoting effect during NeuroD1-mediated AtN reprogramming

The cell survival-promoting effect of overexpressed miR-375 at the early stage of AtN reprogramming could be due to expression inhibition of its target genes. As we have shown that miR-375 can regulate nELAVL levels during neuronal reprogramming (Fig. 4), we decided to examine the roles of nELAVLs in this cell survival effect. We generated retroviral constructs constitutively expressing ELAVL2 and 4 genes without 3’-UTRs (thus miR-375-refractory). The protein expression of these constructs was confirmed by western blot (Fig. 10A). First, when we overexpressed nELAVLs in HA cultures, we found that both nELAVLs constructs induced neurite-like process extension from the infected cells at 6 DPI (Fig. 10B and 10C). However, these nELAVL-infected HA do not express the neuronal markers DCX and NeuN (data not shown) and disappear soon after the morphological change. Indeed, we detected a significantly higher percentage of CCasp3+ cells among the ELAVL4-infected HA compared with control ones at 3 DPI (Fig. 10D), suggesting that overexpression of nELAVLs induced apoptosis in HA. Next, we overexpressed these miR-375-refractory nELAVLs constructs along with NeuroD1-RFP and miR-375. We could not distinguish between nELAVLs and miR-375 infected cells since both expression constructs have a GFP reporter. However, relatively high coinfection efficiency by two retroviruses (Fig. 7B) implied that a significant population was coinfected by three constructs. We then examined CCasp3+ cells among the NeuroD1-infected (RFP+) population and found that additional overexpression of ELAVL4 but not ELAVL2 induced significantly more CCasp3+ cells among RFP+ cells when compared with NeuroD1 and miR-375 coinfection (Fig. 10E). Thus, these data indicate that overexpression of miR-375-refractory ELAVL4 reverses the miR-375-induced survival-promoting effect during AtN reprogramming.

**Figure 10.**
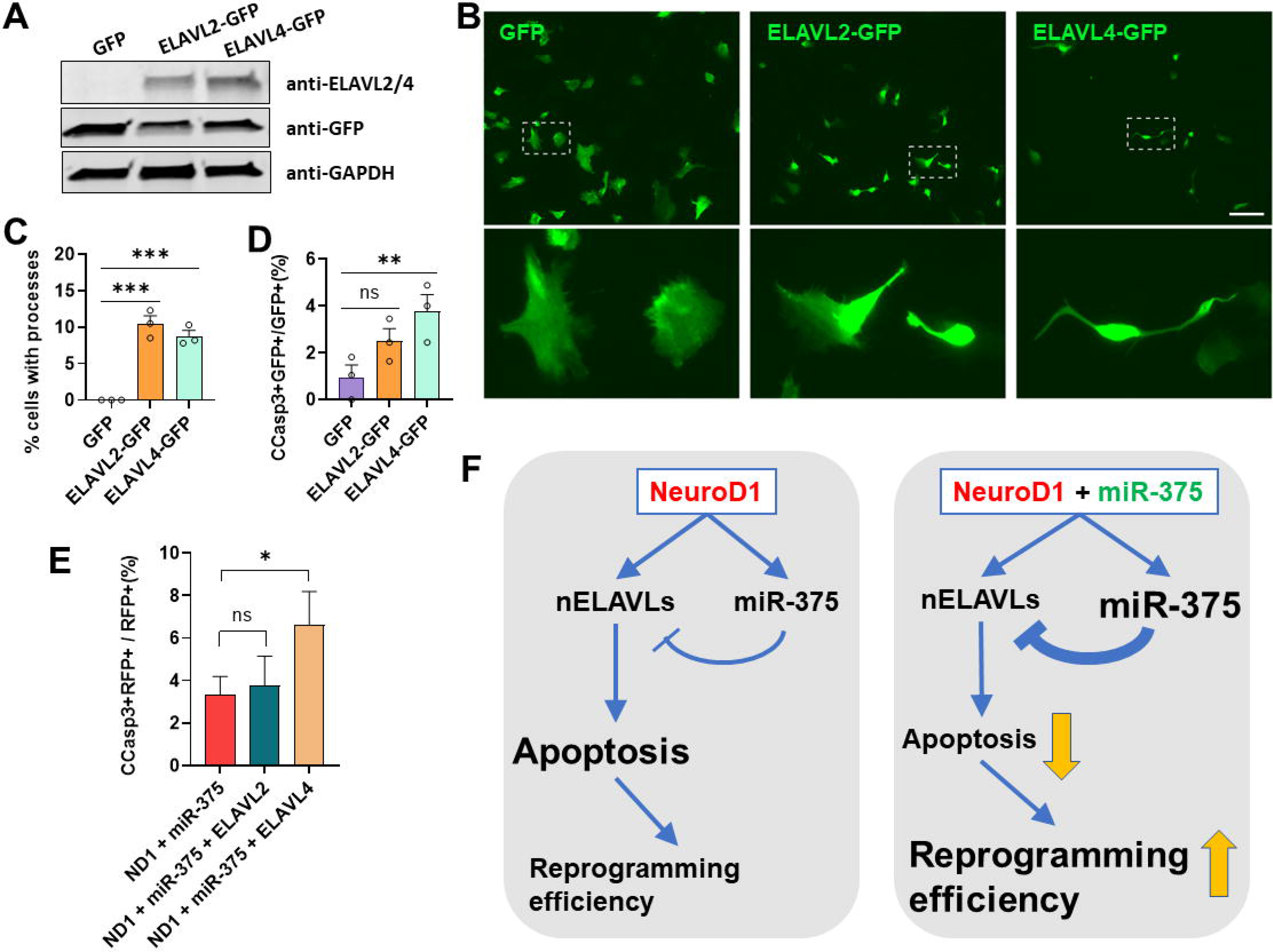
Overexpression of miR-375-refractory nELAVLs induces morphological change and apoptosis in HA, and reverses miR-375-induced survival-promoting effect during NeuroD1-mediated AtN reprogramming. (A) Western blot analysis of ELAVL2 and ELAVL4 protein in retro-ELAVL2-or retro-ELAVL4-infected HA cells at 4 DPI. An antibody anti-ELAVL2/4 was used to recognize both proteins. (B) Representative live florescence images of HA infected with GFP, ELAVL2-GFP, or ELAVL4-GFP retroviruses. Target cells are enlarged to show morphology. Scale bar, 100 μm. (C) Quantitation of the percentage of cells with neurite-like process in HA infected with GFP, ELAVL2-GFP, or ELAVL4-GFP retroviruses. (D) Quantitation of the proportion of CCasp3+ cells in HA infected with GFP, ELAVL2-GFP, or ELAVL4-GFP retroviruses at 3 DPI. (E) Quantitation of the proportion of CCasp3+ cells among RFP+ HA in different coinfection cultures as indicated at 3 DPI. (F) Schematic depicting the normal function of miR-375 during ND1-mediated AtN reprogramming as well as when overexpressed. Data represent mean ± SEM. *, p<0.05; **, p<0.01; ***, p<0.001; ns, not significant.

## DISCUSSION

In this study, we examined both mRNA and miRNA expression changes during NeuroD1-mediated AtN reprogramming by RNA-seq analysis. In addition to significantly upregulated mRNAs related to neuron differentiation, two major upregulated miRNAs, miR-375 and miR-124, were identified. We further investigated the function of miR-375 in reprogramming by overexpression and knockdown approaches and found that miR-375 regulates the expression level of nELAVLs, which are also upregulated by NeuroD1, in the reprogramming astrocytes. Surprisingly, we observed a cell survival-promoting effect by overexpression of miR-375 during reprogramming. Finally, we propose a mechanistic model, in which NeuroD1 turns on miR-375 to modulate its downstream effector genes such as *nELAVLs* to achieve successful reprogramming accompanied by apoptosis, and this miRNA-mediated gene expression modulation can be further tweaked by miR-375 overexpression for a better reprogramming outcome (Fig. 10F). To our knowledge, this is the first study to systematically examine miRNA expression and decipher their functions during glia-to-neuron reprogramming.

### MiR-375 is a transcriptional target of NeuroD1

Our RNA-seq data indicate that miR-375 is one of the most significantly upregulated miRNAs during AtN reprogramming. This result is not completely unexpected since miR-375 has been identified as a direct transcriptional target of NeuroD1 in pancreas (Keller et al., 2007) where it plays a critical role in pancreas development and insulin secretion (Kloosterman et al., 2007; Poy et al., 2004; Poy et al., 2009). However, miR-375 is not well studied in the CNS. It has been reported that miR-375 is transiently expressed in the developing telencephalon at early embryonic stage (E13) before it is downregulated (Abdelmohsen et al., 2010), and yet the regulation of this transient expression of miR-375 is not well understood. The expression level of miR-375 is minimal in the adult CNS except for the pituitary gland (Kapsimali et al., 2007). In a human ES cell culture system, miR-375 has been identified as one of the REST downstream miRNAs, and its expression is upregulated in *REST*-null ES cells and ES-derived neural stem cells (NSCs) (Bhinge et al., 2016). In the same study, the authors have also shown that miR-375 expression increases during motor neuron differentiation from ES cells (Bhinge et al., 2016). To our knowledge, our study is the first to show that miR-375 is upregulated by overexpression of NeuroD1 in neural cells, and this upregulation of miR-375 is likely through direct transcriptional activation by NeuroD1 as is the case in pancreas. Interestingly, miR-124 (another major significantly upregulated miRNA in our RNA-seq analysis) and nELAVLs are also expressed in pancreas, suggesting that common mechanisms of gene regulation and function may exist between CNS and pancreas (Baroukh and Van Obberghen, 2009; Juan-Mateu et al., 2017).

### MiR-375 regulates levels of nELAVLs during AtN reprogramming

The function of nELAVLs in neuronal differentiation during development has been documented. Gene knockout studies have shown that deficits in axonal transport and abnormalities in neuronal polarity are observed in *ELAVL3*-null Purkinje cells in the cerebellum (Ogawa et al., 2018), and that loss of *ELAVL4/HuD* results in a defective dendritic overgrowth in both the neocortex and the CA3 region of the hippocampus (DeBoer et al., 2014). On the other hand, overexpression of *ELAVL4/HuD* accelerates dendritic outgrowth in cortical neuron culture via GAP43 (Anderson et al., 2001). *ELAVL4/HuD* has also been shown to promote neuronal differentiation of neural stem/progenitor cells (NSPCs) in the adult subventricular zone (SVZ) by stabilizing the mRNA of *SATB1*, a critical transcriptional regulator during neurodevelopment (Wang et al., 2015). Furthermore, post-transcriptional regulation of neuronal mRNAs by ELAVL4/HuD has been shown to mediate synaptic plasticity in mature neurons (Tiruchinapalli et al., 2008). With all the above findings, it is not surprising that we observe a drastic upregulation of *nELAVLs* during NeuroD1-mediated AtN reprogramming. We don’t have experimental evidence that *nELAVLs* are direct transcriptional targets of NeuroD1 although it is possible since upregulation of these genes occurs as early as 2 DPI (Fig. 4). Alternatively, this upregulation by NeuroD1 is indirect and occurs via other intermediate transcription factors. Consistently, Neurog2 has been shown to bind two E-boxes in the *ELAVL4/HuD* promoter region and regulate its transcription (Bronicki et al., 2012), and *Neurog2* is also upregulated during NeuroD1-mediated AtN reprogramming (data not shown).

*ELAVL4/HuD* is a validated target gene of miR-375, overexpression of which reduces *ELAVL4/HuD* gene expression and in turn results in decreased dendritic branching in the mouse hippocampus (Abdelmohsen et al., 2010). MiR-375 is transiently expressed in the neocortex during embryonic development, and its temporal expression pattern is complementary to that of *ELAVL4/HuD* suggesting a negative regulatory control between miR-375 and *ELAVL4/HuD* (Abdelmohsen et al., 2010). Our data in this study further confirm that miR-375 can regulate nELAVLs during neuronal reprogramming process especially at early stages (Fig. 4). We postulate that the significance of this miR-375-mediated regulation is to tune NeuroD1-induced nELAVLs expression to an optimal level for successful reprogramming. Consistently, it has been reported that although *ELAVL4/HuD* expression is increased after learning and memory, constitutive *ELAVL4/HuD* overexpression in the adult brain does not improve, but rather impairs animal behavior in a battery of learning and memory tests (Bolognani et al., 2007), indicating that tightly controlled ELAVL4/HuD expression level is critical for its function.

Neuronal reprogramming is not a “natural” biological process (Chen and Li, 2022), and constitutive overexpression of reprogramming factors may trigger distinct cellular and molecular responses in the reprogrammed cells. In our case, forced expression of NeuroD1 in astrocytes initiates neuronal reprogramming by turning on downstream genes including *nELAVLs*. However, high level of *nELAVLs* expression not only facilitates reprogramming process, but also brings detrimental effects such as apoptosis. The NeuroD1-induced expression of miR-375 is probably an adaptive response of the reprogrammed cells to trim down *nELAVLs* expression level for a better outcome. We propose that this “incoherent” (Shkumatava et al., 2009) regulation between miRNAs and their target genes is not a rare event during NeuroD1-mediated reprogramming since we observed many miRNA/target gene pairs that are both upregulated by NeuroD1 (Fig. 3). Furthermore, this expression “trimming” function of miRNAs may be common in many transcription factor-induced reprogramming scenarios. Another point worth noting is that *nELAVLs* are also upregulated in other glial cell types including microglial and glioblastoma cell lines during NeuroD1-mediated reprogramming (Matsuda et al., 2019; Wang et al., 2021b). It would be interesting to examine if miR-375 is also upregulated in these glial cells and modulates *nELAVLs* expression level during these neuronal reprogramming processes.

### MiR-375 is neuroprotective during AtN reprogramming

AtN reprogramming is an event involving drastic changes in gene expression and signaling pathways. Many cells die during the reprogramming process if they cannot pass the metabolic “checkpoint” (Gascon et al., 2016). Cells that manage to survive will proceed to convert into neurons. Our previous research has indicated that NeuroD1 can convert reactive astrocytes into functional neurons in the spinal cord with high efficiency (Puls et al., 2020). This high efficiency is partially due to the fact that NeuroD1 is not only a critical neuronal differentiation transcription factor, but also a survival factor during development, especially in developing granule neurons of the cerebellum and hippocampus where NeuroD1 is highly expressed throughout adulthood in mice (Miyata et al., 1999). Despite this fact, we did observe cell apoptosis at the early stage (3 DPI) of reprogramming in cultured HA after NeuroD1 overexpression and surprisingly miR-375 help reduce apoptosis when co-expressed with NeuroD1 (Fig. 8). In fact, the survival-promoting effect of miR-375 has been documented previously. For instance, the miR-375 level is reduced in the degenerating motor neurons of the ALS mouse and human patients (Bhinge et al., 2016; De Santis et al., 2017), and knockdown of miR-375 in culture induces motor neuron death via p53 pathway (Bhinge et al., 2016).

Surprisingly, we did not detect any functional effects of inhibitor-mediated miR-375 knockdown during reprogramming process in HA cultures except for an increase of ELAVL2/4 protein levels (Fig. 4F). This could be due to only partial knockdown of miR-375, and thus miR-375 gene knockout may help reveal potential loss-of-function effects if any. Nevertheless, our data show that high levels of nELAVLs (i.e. by overexpression) induces apoptosis in HA culture (Fig. 10) and miR-375 can regulate *nELAVLs* expression level during NeuroD1-mediated reprogramming (Fig. 4). Therefore, it is likely that miR-375 helps suppress NeuroD1-induced *nELAVLs* expression to elicit cell survival-promoting effect (Fig. 10F). However, we don’t exclude other pathways (such as p53) that may also be affected by miR-375 overexpression (Bhinge et al., 2016) and are independent of nELAVLs. Regardless of the potential mechanisms, our observation that miR-375 is neuroprotective during neuronal reprogramming has translational values. Overexpressing miR-375 along with NeuroD1 could further elevate cellular miR-375 level (Fig. 7H) and increase neuronal reprogramming efficiency by promoting survival of reprogrammed neurons. We are currently constructing the combinatory vector to co-express NeuroD1 and miR-375, and the reprogramming efficiency of this co-expression vector will be characterized in our future investigation.

## ACKNOWLEDGEMENTS

This work was supported by startup funds from Medical College of Georgia at Augusta University (to HL), National Institutes of Health grants (R01NS117918, R21NS104394, R21NS119732) (to HL) and T32GM108563 (to IS), and Ann L. Jones Spinal Cord Regeneration Research Fund (to HL).

## AUTHOR CONTRIBUTIONS

H.L. and X.C. conceived the idea and supervised the entire project. X.C. performed the major experiments, analyzed the data, and made the figures. X.W., M.J., N.M., C.W., N.J., and K.M. contributed to the experiments. I.S., B.P., and Y.S. helped with RNA-seq analysis. Q.D. provided expertise and guidance on viral constructs and packaging. H.L. wrote the manuscript and secured funding.

## DATA AND MATERIALS AVAILABILITY

The RNA-seq datasets generated in this study have been deposited in GEO, GSE214749. The raw data supporting the conclusions of this article will be made available by the authors, without undue reservation.

